# An information sign changes the way the public perceives exotic pond sliders *Trachemys scripta* in the Altrhein of Kehl (Germany)

**DOI:** 10.1101/2022.07.19.500557

**Authors:** Carsten Schradin

## Abstract

Invasive pond sliders (*Trachemys scripta*) have been released in thousands of freshwaters within Europe and reproduce in the southern states and even in warm areas of Germany. All member states of the European Union must have an action plan how to fight this invasive species. The German action plan focusses on informing the public, but to date no study investigated the impact of such actions. Instead, NGOs setting up information signs report that this does not stop the release of exotic pond turtles, but without any quantitative measure. In 2019, we put up an information sign at the Althrein of Kehl, an oxbow lake where for the first time it had been shown that *T. scripta* is breeding in Germany. I performed interviews with people walking along the oxbow lake before the sign was pit up in 2019, and again in 2022. Counts of exotic pond turtles still increased, but this was mainly due to an increased number of small pond turtles, while numbers of very large ones did not further increase. This indicates that the increase in peak counts might be rather due to local reproduction than additional release. After the information sign was set up, more people responded that the presence of exotic pond turtles is problematic for nature conservation and animal welfare, that it’s illegal to release them, and that they should be removed. This response was especially strong in people who had read the information sign. Independent of the information sign, most interviewed people stated that one should not release pond turtles into the wild, but bring them to animal shelters. While the data here only represent one single case study that might not be representative, it’s the first study showing that putting up information signs is effective in changing the attitude of people who had read it. This indicates that investment into informing the public is worthwhile, but also that at the same time evaluations of the impact of the measures are important. National action plans should focus on a combination of informing the public and removing the exotic pond turtles, but also on providing keepers of these animals the option to leave the animals at an animal shelter instead of releasing them into the wild.

## INTRODUCTION

Invasive species, i.e. exotic species introduced by humans that establish themselves outside their natural distribution range, are threatening native biodiversity worldwide (Geiger & Waitzmann 1996; Wilson *et al*. 2009). One reptile species that has become invasive in Central and South America, Africa, Asia, and Europe is the north American red-eared slider (*Trachemys scripta*) (Böhm 2013; Standfuss *et al*. 2016; Mo 2019). This species is also widely distributed in Europe, where it has been released by pet owners into thousands of fresh waters (Cadi *et al*. 2004; Prevot *et al*. 2007; Kopecký, Kalous & Patoka 2013; Standfuss *et al*. 2016).

The European Union identified *T. scripta* as an invasive species (European_Commission 2016) against which the member states must take action to prohibit the import, breeding and release of this species (European_Parliament 2014). As environmental conditions differ between member states, the national action plans also differ. In south European countries like Spain and France, where this species breeds and spreads very fast, removal of exotic pond turtles is one main action. In Germany, the main proposed action against *T. scripta* is to increase public awareness (StA_„Arten-_und_Biotopschutz” 2018), which is also part of the general actions proposed by the European Union (European_Parliament 2014). However, I am not aware of any study investigating the effects of such actions. While signs not to release exotic pond turtles have been set up at some localities, for example in Munich by the Reptilienauffangstation (https://www.reptilienauffangstation.de), effects of these public awareness actions have not been measured. Instead, as the release of exotic pond turtles continued seems to have continued, frustration about the low effectiveness has been high (anonymous communication by different NGOs). However, it was never measured, only assumed, that release continued and the public awareness actions were ineffective. So far it is unknown whether such information signs influence the awareness of the public and lead to a decrease in the release of exotic pond turtles.

Here I present a case study conducted at a oxbow lake, the Altrhein of Kehl, the only location in Germany for which so far successful reproduction of *T. scripta* has been reported (Schradin 2020). This population has been monitored by me since 2016. In 2019, I set up an information sign for the public and I continued monitoring the population, getting an estimate whether it was still growing. Before the information sign was set up, I conducted interviews with people walking along the oxbow lake, asking them about how they evaluate the presence of exotic pond turtles. These interviews were repeated in 2022. If the information sign had a positive impact in educating the public, I predicted that in 2022 1. more people to regard the presence of exotic pond turtles to be problematic, 2. because they know it poses problems for both nature conservation and animal welfare, 3. which is why the release of exotic pond turtles is illegal, 4. and instead they should not be released, 5. but removed from the oxbow lake.

## MATERIALS AND METHODS

### Study area and study period

The study was conducted from 2016 to 2022 at the Altrhein of the city Kehl (48° 34′1.95″N, 7° 48′35.41″E), which is an 90m long and 25 to 80m wide oxbow lake formed over 100 years ago from the River Rhine. Kehl is in the Upper Rhine Valley, the warmest area of Germany. A community of six different species of exotic pond turtles exists in the Altrhein, which has been continuously growing in population size from 2016 to 2020 (Schradin 2020). Of these, *T. scripta* is the most common species, and both clutches and hatchlings have been found in several years, proofing for the first time successful reproduction of this species in Germany (Schradin 2020).

### Monitoring

In the years 2016-2020, the population of exotic pond turtles was monitored on six to eight afternoons per year, during April to July. These data are already reported in a previous publication (Schradin 2020), but added here for the question whether there was indication of further release of exotic pond turtles after the information sign was put up. Due to the Corona lockdown in 2021, and the concentration on getting more data from interviews in 2022, during these two years turtles were counted only during 4 afternoons per year. In all cases, observations were made using binoculars at five locations along the eastern shore of the lake, that were previously determined to have a high abundance of pond turtles. In addition, any pond turtle observed between these locations was recorded. For every individual, the carapace length was estimated to be in one of the following categories: 5cm (hatchlings), 10cm, 20cm or 30cm.

### Setting up an information sign

An information sign was created by me during a course I give at the Hector Akademie Kehl. In this course, I teach highly gifted school children 8-9 years old about ecology and nature conservation, with the exotic pond turtles being an example. The information sign and its translation into English are shown in Figure 1. The sign informs about where the animals come from, how many species there are, that one species is invasive, and why releasing exotic pond turtles is a problem for nature conservation and animal welfare, and thus illegal. As the German city of Kehl is next to the French city of Strasbourg and visited by many French citizen, with several thousand French citizens living in Kehl, the information sign also includes a summary in French (Fig. 1).

**Figure 1.**
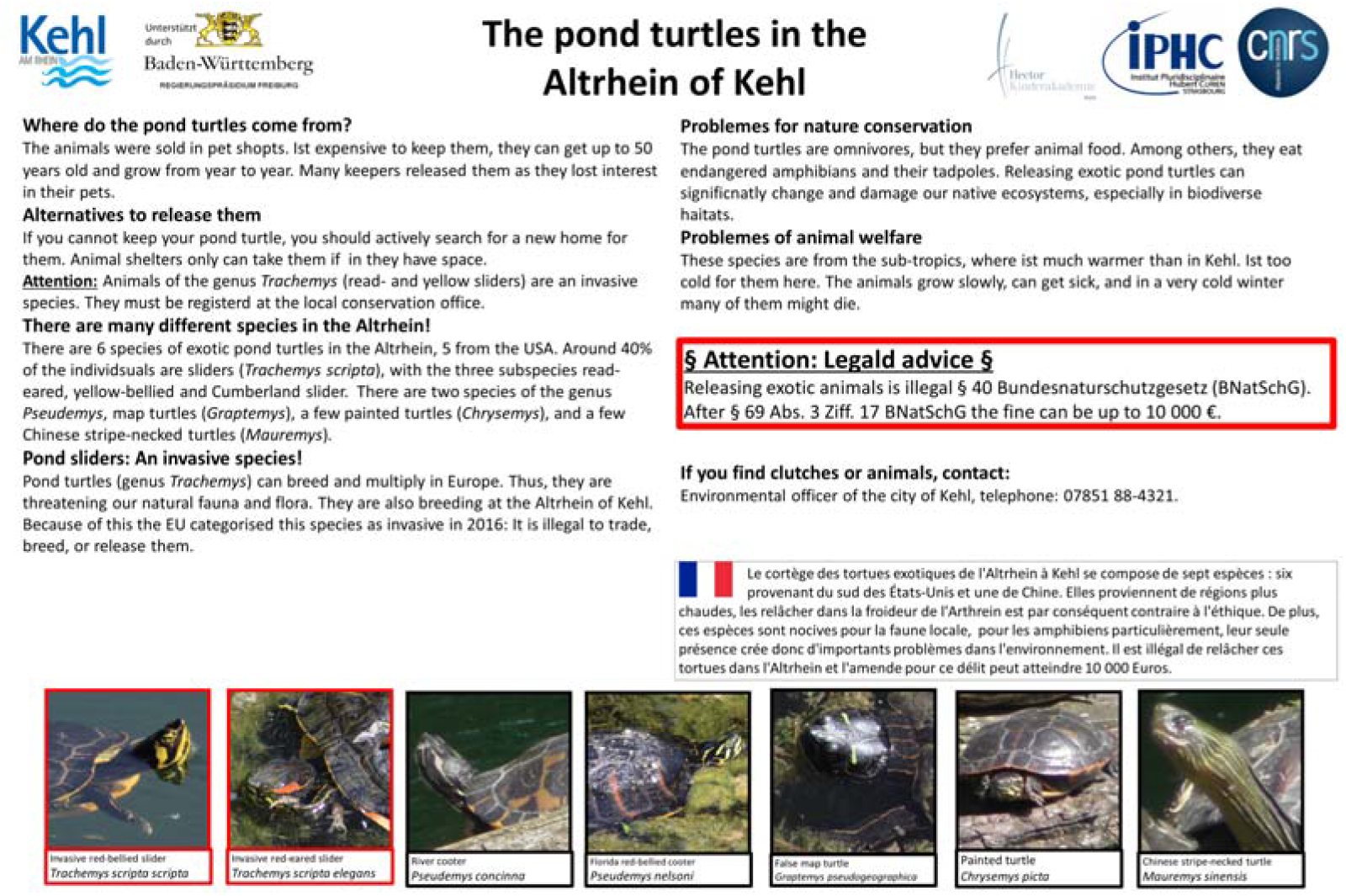
Top: The information sign put up in July 2019.

The city of Kehl installed the information sign at a bridge over the Altrhein next to the communal hospital on the 16^th^ of July 2019. This spot was chosen as from here people often observe the exotic pond turtles sun-basking on some branches of a dead tree that are lying in the water.

### Interviews

Interviews were conducted during 5 afternoons in June and July 2019 and 4 afternoons in May and June 2022. In 2019, 28 people were interviewed, in 2022 30 people, 13 of which had read the information sing. Interviews were conducted by myself and by pupils from the Hector course I was teaching. The pupils were well instructed and the first interview they watched me and I was present while they performed interviews. As the interviews were anonymous, no ethical clearance was needed after French law and CNRS administration.

First the pupils introduced themselves, that they are from the Hector Kinderakademie and do a survey about the exotic pond turtles. They made clear its no test but the aim is to find out what people know about these animals. We only used the term “pond turtles”, without “exotic” during the interviews. The questions were identical in both years. Only in 2022 we added as very last the question whether the interviewee had read the information sign or not. Only after the interview could the interviewees ask questions and get more information, if they wanted.

The data sheet for the interviews included for the pupils how to classify the answers, and they had been trained to do so. The questions and the possible categorised answers (in brackets) were:

1. Do you find it problematic that there are pond turtles in the Altrhein? (yes / no).
2. Why do you think its problematic? (not problematic / animal welfare / nature conservation / other / don’t know).
3. Is it legal to release pond turtles here? (is illegal / is legal / don’t know).
4. In your opinion, what should someone do who has a pond eared turtle as pet but can no longer keep it, e.g. because they no longer have time or are moving away? (animal shelter / sell/ keep/ don’t know).
5. What should happen to the pond turtles in the Altrhein? (leave them and don’t disturb / trap and remove / kill / don’t know).

### Data analysis

Data on pond turtle abundance are expressed as peak counts, the maximum number of live individuals observed in one survey afternoon of a particular year. The data from the years 2016-2020 have already been published (Schradin 2020), but are included here for comparison. Data from the interviews were analysed by comparing the ratios of correct answers vs. unknown plus wrong answers, using the Fisher’s Exact test.

## RESULTS

### Turtle numbers

The annual total peak counts increased continuously from 2016 to 2020, even after the information sign was put up (Fig. 2). After the count in 2020 (Mai-July), 58 individuals were trapped and removed end of July 2020. This influenced the expected pond turtle numbers for the next year: if there was no recruitment in turtle numbers, I expected for 2021 the peak count from 2020 minus 58; instead, it was higher in 2021 and further increased in 2022 (Fig. 2). The number of very large pond turtles (carapax size 30cm) did not increase after the information sign was put up and was not higher than the expected peak number. However, the number of large pond turtles (carapax size 20cm) increased, which could be due to medium sized pond turtles (carapax size 10cm) growing, as their numbers decreased (Fig. 2).

**Figure 2.**
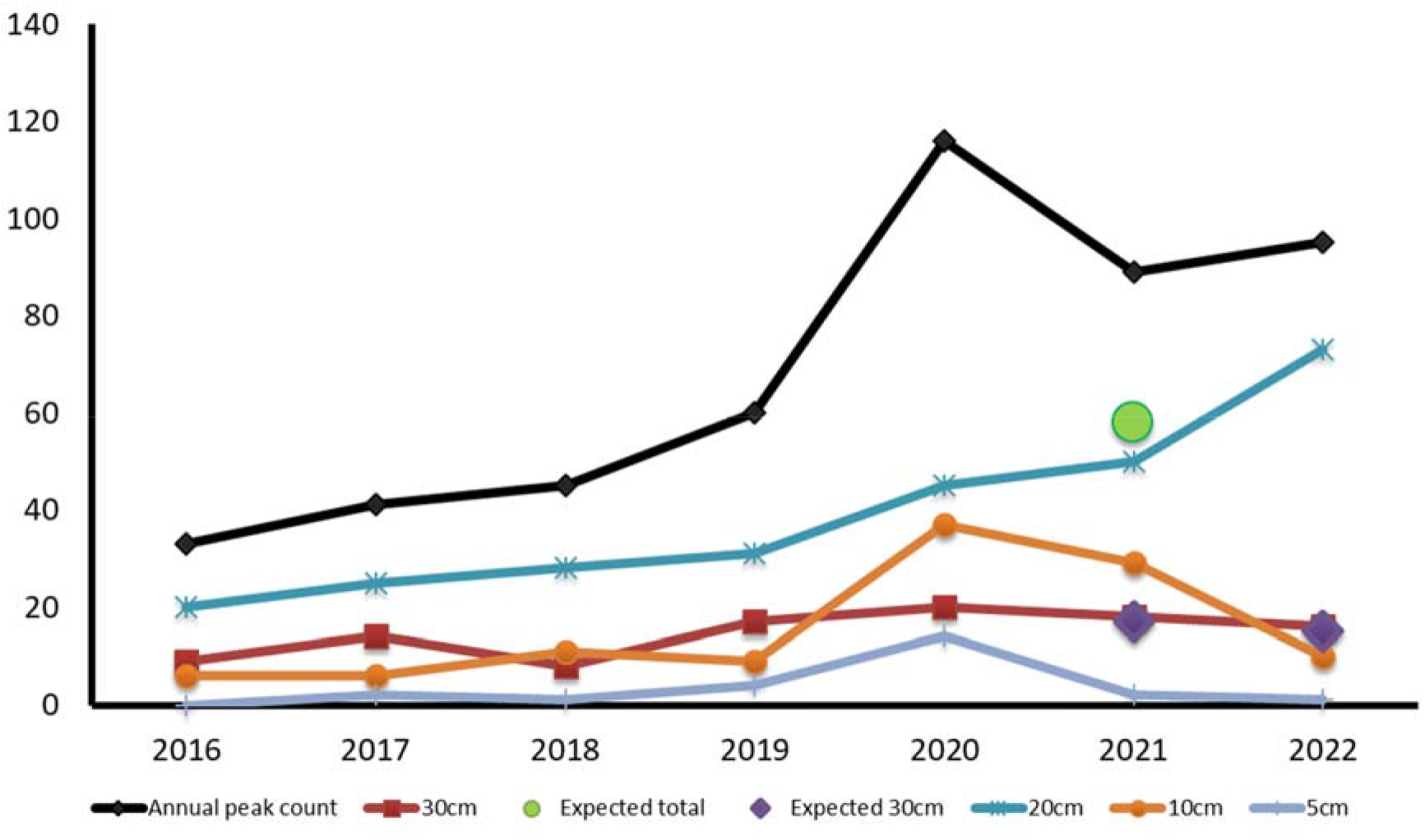
Annual peak count for all turtle species in the Althrein of Kehl. In black, total numbers. In red shown are the very large turtles which are most likely to be released, in other colours smaller turtles. After the count in 2020, a total of 58 turtles were removed, indicated by the green spot as expected value for 2021 (count 2020-58). Of these 58 turtles, one was very large; additionally, two large turtles caught on land were removed in 2021. The diamonds indicate the number of expected large turtles if no new ones were released.

### Responses during the interviews

There was no statistical difference in the proportion of people reporting the pond turtles to be a problem between 2019 and 2020 (p=0.28; Fig. 3). However, when only considering the people that had read the sign, then significantly more regarded the pond turtles as problematic, both compared to 2019 (p=0.02) and to the people in 2022 who had not read the sign (p=0.02; Fig. 3).

**Figure 3.**
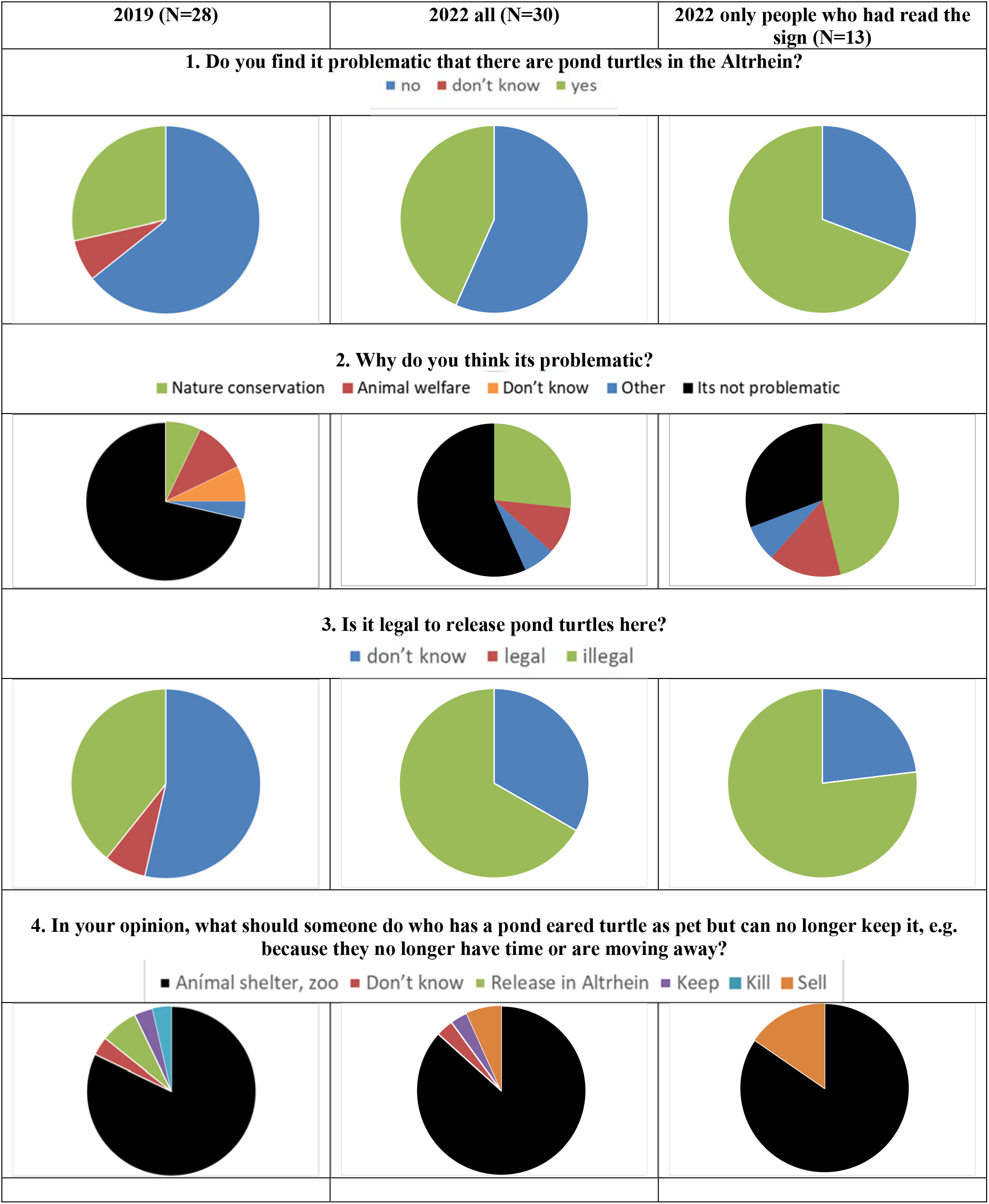
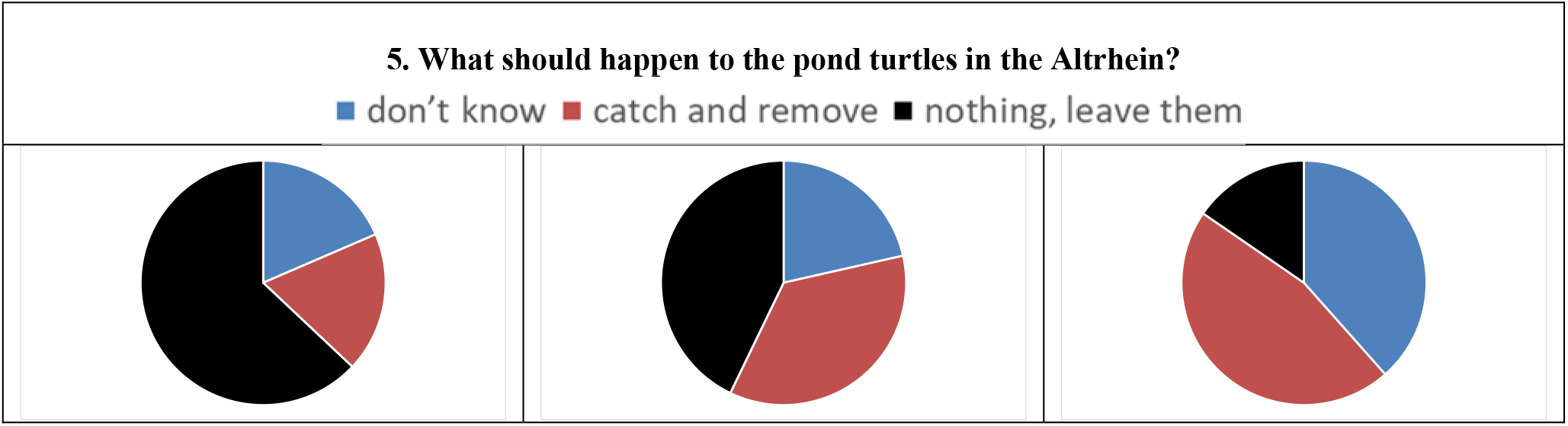
Responses to specific questions of the interview, in 2019 (left; before the information sign was put up), in 2022 (middle; after the sign was in place for 2), and in 2022 by respondents who said they had read the information sign (right).

In 2022, not more people reported nature conservation and /or animal welfare to be a problem than in 2019 (p=0.15; Fig. 3). However, when only considering the people that had read the sign, then significantly more identified nature conservation and /or animal welfare to be the problem, both compared to 2019 (p=0.02) and to the people in 2022 who had not read the sign (p=0.01; Fig. 3).

In 2022, more people assumed it to be illegal to release pond turtles than in 2019, though the difference was not significant (p=0.06; Fig. 3). Considering only the people that had read the sign, the difference was significant (p=0.04; Fig. 3), while the difference between people who had and who had not read the sign in 2022 was not significant (p=0.44, Fig. 3).

Asked what somebody should do with a pond turtle he has as pet if he cannot keep it anymore, then in both years most people suggested that pet owner should bring the pond turtle to an animal shelter or a zoo if for whatever reason they cannot keep it anymore (Fig. 3). Only in 2019 did two people suggest to release them in the Altrhein (Fig. 3).

In 2022, not more people suggested to trap and remove the exotic pond turtles from the Altrhein than in 2019 (p=0.23; Fig. 3). Considering only the people who had read the sign, the difference was more pronounced, but did not reach significance (p=0.07, Fig. 3).

## DISCUSSION

In this case study, I found that the awareness of the general public for the problem of invasive *T. scripta* increased significantly two years after an information sign was put up. At the same time, there was no evidence that a large number of additional pond turtles were released,. Therefore, this study gives the first empirical support that informing the public is a suitable tool of the action plan against invasive exotic pond turtles

The current study has several shortcomings reducing its general significance. First, it’s a minimal sample size of one sign at one location. In how far similar results would be found at different localities and different information signs is unknown. But this study indicates that it is worthwhile to try informing the public using such signs and then to evaluate whether it caused increased awareness. Second, the monitoring of the population was done without individual identification (like capture mark recapture), and during very few afternoons. As such, the data do not allow me to conclude that no or only few pond turtles were released after the sign was put up. Nevertheless, the data do not provide evidence that a large number of additional exotic pond turtles have been released. If the pond turtles could be identified individually, for example via photos and the use of AI, it would be possible to identify which animals are recruited from year to year into the population, and whether they are small (possibly due to reproduction) or large (possible releases).

Comparing responses to the interview questions before and two years after the information sign was put up indicated a clear change in publica awareness: After the information sign was set up, many more people were aware that the exotic pond turtles represent a problem for nature conservation and / or animal welfare, that its illegal to release them, and that it would be appropriate to remove them. Theoretically, this response could have been influenced by many co-factors and not the information sign alone, for example by reports in the local newspaper about the problem, or by people having had more time to walk along the Altrhein during the Corona pandemic and then inform themselves at home about the exotic pond turtles they saw. However, the difference in awareness was most obvious in people that reported to have read the information sign, and these were also significantly better informed than people interviewed during the same period in 2022, that had no read it. Thus, the most parsimonious explanation is that reading the information sign increased the awareness about the problem of exotic pond turtles.

The answer to one question did not differ between years and was not dependent on whether or not the interviewees had read the information sign: What should somebody do with his exotic pond turtle pet if he cannot keep it anymore, for example because he is moving. Nearly nobody suggested to release the pet into the Altrhein, but the large majority suggested to bring them to a zoo or an animal shelter. The problem is that zoos are not interested in taking exotic pond turtles, nor are animal shelters, most of which don’t have the facility for this nor the funding. This can explain why ten thousands of these pets have been released within Europe, as the alternative to give them to an animal shelter does not exist. In reverse this means that if we want to avoid more exotic pond turtles to be released, then we must create this opportunity, by providing funding to animal shelters. In Germany, private organisations exist, like the Reptilienauffangstation in Munich (https://www.reptilienauffangstation.de), but these are heavily underfunded. National and regional authorities interested in reducing the number of releases should provide funding for such organisations, and funding to local animal shelters to provide facilities to keep exotic pond turtles.

The European Union demands that all member states should take action against invasive species (European_Parliament 2014) including *T. scripta* (European_Commission 2016). In Germany, it has been believed that the it is too cold for the released individuals to survive for long periods or to reproduce (Geiger & Waitzmann 1996; Pieh & Laufer 2006; Laufer 2007; Nehring 2016). The German action plan focusses on increasing public awareness (StA_„Arten-_und_Biotopschutz” 2018), but this is neither enforced nor evaluated. A previous study proofing successful reproduction of *T. scripta* at the Altrhein of Kehl (Schradin 2020) and the ever increasing population size there (Fig. 2) make it clear that public awareness has to be increased, and that additional actions are required (Teillac-Deschamps *et al*. 2009).

In conclusion, the information sign was successful in increasing public awareness. The national action plan (for Germany and all other countries) for *T. scripta* should include three main components: 1. Increasing public awareness by providing information, and evaluating the effectiveness of these actions. 2. Providing funding to animal shelters to take in exotic pond turtles. 3. Removing *T. scripta* from natural habitats to avoid them establishing viable and spreading populations (Cadi *et al*. 2004; Sancho & Lacomba 2016).

## ACKNOWLEDGEMENTS

This project was funded by the CNRS. The project was done as part of a course I do at the Hector Kinderakademie in Kehl, teaching 8 to 9 years old children about animal behaviour, population biology, animal welfare and nature conservation. I am very thankful to the Hector Kinderakademie in Kehl for their support and to the kids who conducted interviews during the two years: in 2019 Tabea Dedic, Celine Elysev, Lucy Fleig, Theo, Kreipl, Alena Lorenz, Emma Sauer, and Sebastin Sirbu. In 2022 Ellen Bähr, Marie Chambrion, Lena Hummel, Elena Humpfer, Emilia Reichmann, Paul Sauer, Ivy Senn and Fabian Speck.

